# Levodopa increases substantia nigra iron: implications for Parkinson’s disease

**DOI:** 10.64898/2026.05.26.728066

**Authors:** Guangwei Du, Luke Bransom, Mi Zhou, Christopher Sica, Xuemei Huang, Yang Yang, Richard B. Mailman

## Abstract

**Background:** Excessive or unregulated iron in the brain can lead to toxicity via ferroptosis and related mechanisms. Iron accumulation in the substantia nigra (SN) occurs with Parkinson’s disease (PD) progression and has been hypothesized to be an etiological mechanism.

**Objective:** Based on emerging clinical observations, we tested the hypothesis that iron accumulation in the SN is a consequence of levodopa administration and is treatment-related rather than an intrinsic etiological mechanism.

**Methods:** We used both unilaterally lesioned 6-OHDA and unlesioned rats. We administered levodopa to rats at doses that were allometrically calculated to be similar to those used in mid-stages of PD. Iron-sensitive MRI (R2*) was used to quantify iron in the brain. Both group and intra-subject analyses were done using paired t-tests and linear mixed models.

**Results:** Experiment 1 used the unilateral 6-OHDA model to take advantage of the almost complete lack of dopamine neurons on the lesioned side. This permitted testing if levodopa-induced iron accumulation occurred in and/or depended on dopamine neurons. Fifteen days of levodopa treatment caused a marked increase in Fe in both the lesioned (p = 0.042) and unlesioned sides (p = 0.005), showing that iron accumulation does not depend on the presence of dopamine neurons. Based on these data, in experiment 2 unlesioned rats were administered levodopa daily for four months, and iron (R2*) values were assessed at baseline, 1, 2, and 4 months. In these normal rats, the levodopa-treated group had significantly increased Fe (R2*) in the substantia nigra compared to the vehicle group (p = 0.013). Interestingly, these effects were limited to the striatum, with no increases seen in the striatum, ventral tegmental area, or frontal cortex

**Conclusion:** Levodopa triggers processes that increase iron deposition in the substantia nigra, but this process may not depend on dopamine neurons. The underlying mechanisms and the effect on PD progression are important to elucidate and may transform how we understand PD and related neurodegenerative disorders

## Introduction

Parkinson’s disease (PD) is marked pathologically by loss of dopamine neurons in the substantia nigra (SN) of the basal ganglia.^1, 2^ Increased iron content at the SN also has been reported by post-mortem studies of levodopa treated PD patients.^3-7^ With the advance of MRI technology, studies using in vivo iron sensitive MRI demonstrated that the increased SN iron content can be seen even in the early stage of disease.^8^ The exact mechanism leading to increased iron in the SN, is unclear, nor is it known if it is related to disease progression and/or antiparkinsonian medication.^9^

Iron is an essence trace metal involved in many important neurobiological processes, including myelin production, neurotransmitter synthesis, mitochondrial function, etc.^10^ Findings from preclinical studies have suggested that excess intracellular iron is associated with increased reactive oxygen species and oxidative stress, cellular/mitochondria dysfunction, and neuronal death in PD, possibly playing a role in PD pathogenesis.^11-13^ Importantly, iron and dopamine can form an iron-dopamine complex that has been proposed as a neurotoxic mechanism that might lead to dopamine neuron loss in PD.^14-17^

Levodopa is the current mainstay of antiparkinsonian drugs because it can restore lost dopaminergic tone and alleviate PD symptoms effectively. As the precursor of dopamine, excessive levodopa might interplay with iron and lead to oxidative stress and cascading neuron loss.^18^ This notion, however, has been outweighed by benefits of its outstanding symptom control even if there is accelerated loss of dopaminergic terminals (e.g., in the ELLDOPA trial).^19, 20^ The potential importance of iron in PD pathogenesis and the necessary use of levodopa in PD therapy warrant a clearer understanding of the underlying mechanisms. Using iron-sensitive MRI, our recent study of a subjects with PD disease of varying stages (from drug naïve to late stage) demonstrated that the increases in nigral iron occurred only after medical treatment began, and thus may be associated with levodopa treatment.^21^ Using a protein-wide target identification technique, our group has predicted that levodopa might interact with iron regulation/transportation pathways, and potentially lead to iron accumulation and toxicity in PD.^22^

Our clinical observation^21^ led to the hypothesis that levodopa was responsible for iron increases in the SN. We used *in vivo* iron-sensitive MRI in two experiments. In the experiment 1, we used the unilaterally lesioned 6-hydroxydopamine (6-OHDA) rat PD model with 15 days of levodopa administration. Based on the resulting data, in experiment 2, we compared the effect of levodopa and vehicles in healthy rats after 4 months of levodopa treatment.

## Methods

### Subjects

Adult male Fischer 344 rats with an average weight of 250-300 grams at arrival (i.e., ca. 3-4 months old) were purchased from Charles River (Wilmington MA), Envigo (Frederick, MD), or NIA (Hollister, CA; Raleigh, NC; or Kingston, NY). They were housed individually or grouped based on their weight and maintained on a 12-hour light-dark cycle with food and water continuously available. All animal care and surgical procedures were in accordance with the National Institutes of Health “Guide for the Care and Use of Laboratory Animals” and with the Penn State College of Medicine Animal Resources Program. The protocol was reviewed and approved by the Penn State College of Medicine Institutional Animal Care and Use Committee.

Our power analysis for the first experiment was based on clinical data showing that iron increases in the SN were significant even after two years when levodopa doses were low. Considering other factors (stage of disease and levodopa dose, low variability in rat subject population versus humans, each animal serving as its own reference, etc.) led to prediction of a large effect size and N = 9 (each animal l). We started with 12 rats to allow for loss of subjects due to bad MRI scans, illness, etc.

### Materials

L-3,4-dihydroxyphenylalanine methyl ester hydrochloride was purchased from Sigma - Aldrich (Saint Louis, MO). The peripheral aromatic amino acid decarboxylase inhibitor benserazide and the dopamine agonist apomorphine were purchased from Tocris (Bristol, UK).

### Drug administration

We chose levodopa dosing based on the following considerations. A high but common clinical dose in PD patients is ca. 600 mg three times a day, or 1,800 mg (as levodopa). For a 60 kg person, this is a daily dose of 30 mg/kg. Using an accepted allometric conversion of 5 for human to rat,^23^ this yielded a rat dose of 150 mg/kg. We therefore chose a dose of 100 mg/kg/day that is significantly lower than a typical high dose, but somewhat adjusts for the fact that we administered the drug in a single bolus as opposed to 3-5 doses that humans would receive in mid-to late-stage PD. The benserazide was administered simultaneously at a dose of 15 mg/kg to inhibit peripheral amino acid decarboxylase.

The first experiment was powered based on adjusted clinical findings to determine if our power estimates were realistic. When a significant effect was found, the second experiment sought to provide a more realistic model of long-term human exposure and also be an independent validation of the first experiment. The length of treatment for experiment 2 was based on our previous work in humans that showed that iron accumulation is clearly evident after ca. 5 years (or less) of levodopa exposure in PD. Assuming a human life span of ∼80 years, we arbitrarily chose to expose rats to levodopa for 15 days, 1, 2, and 4 months, the latter approaching 15% of the mature rat life span.

### Overall experimental design

#### Experiment 1

Twelve rats were unilaterally lesioned with 6-hydroxydopamine (6-OHDA). Four lesioned rats were purchased from Charles River (Surgery Code: PARKINSON)^24, 25^ and eight rats had surgery in-house following the same protocol. After recovery from the surgery, rats were challenged with apomorphine. Lesion effectiveness (i.e., unilateral dopamine neuron loss) was assessed by vigorous contralateral rotations.^26^ The first (i.e., baseline) MRI scan was performed at least nine days after the 6-OHDA surgery. After the baseline scan, a chronic, daily subcutaneous levodopa/benserazide treatment began the following day and continued for 15 days. After the last treatment, a follow-up MRI scan was performed. Iron content of the SN was extracted from susceptibility MRI images and compared between baseline and follow-up for the 6-OHDA lesioned side and unlesioned side separately. The caudate/putamen complex was chosen as a negative control region. One of the subjects had an MRI in the post-treatment session that had poor image quality due to surface coil dysfunction. This subject was excluded from analysis.

#### Experiment 2

To gauge the long-term effect of levodopa induced iron accumulation at the SN without 6-OHDA lesioning, 10 rats were randomly assigned to two groups (treatment group and vehicle group, 5 rats each group). The treatment group received four months of chronic daily subcutaneous levodopa and benserazide injections as in Experiment 1. The vehicle group received four months of daily subcutaneous injection of 0.1% ascorbic acid injections and served as the control group. Four susceptibility MRI scans were obtained at baseline, 1 month, 2 months, and 4 months timepoints.

### MRI data acquisition

All MRI data were collected on a Bruker BioSpin 7T MRI system with a ParaVision 360 version 2.0 console. A T1-weighted image was obtained as anatomical reference using a Rapid Acquisition Relaxation Enhanced sequence with the parameters: repetition time (TR) = 4800 ms; echo time (TE) = 6.5 ms; matrix size = 200 × 200; field of view (FoV) = 35 × 35 mm; slice number = 50; slice thickness = 0.5 mm; voxel size = 0.175 × 0.175 × 0.5 mm^3^. Apparent transverse relaxation images (R2*) was obtained as brain iron measurement using a 3D multiple gradient echo sequence with the parameters: repetition time (TR) = 50 ms; echo time (TE) = 4 to 33.4 ms with a 4.2 ms echo spacing, resulting 8 echoes in total; matrix size = 192 × 192 × 30; field of view (FoV) = 35 × 35 × 22.5 mm; resulting in a voxel size = 0.182 × 0.182 × 0.75 mm^3^. R2* images were estimated by nonlinear curve fitting of a mono-exponential equation 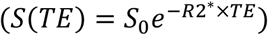, using the Levenberg–Marquardt optimization algorithm.

### Image Analysis

The region of interest (ROI) is defined referring to publicly available 3D MRI Waxholm Space atlas of the rat brain.^27, 28^ The SN was the primary ROI for this study. The ventral tegmental area (VTA), caudate/putamen complex (ST), and frontal lobe (FL) were chosen as self-contrasting regions due to their connections with the SN. The original SN definition was from the Waxholm Space atlas of T2*-weighted template image (Figure 1A). The SN is located at the ventral midbrain area with an oval shape region and an estimated size of 3 × 1 mm^2^ in the Waxholm Space atlas. To avoid inclusion of non-substantia nigra and account for individual anatomical variation, a morphological erosion operation was applied to the SN ROI in the atlas space to obtain a slightly (one voxel) smaller oval shape ROI. In addition to the SN, a round shape ROI (0.8 × 0.8 mm^2^) also was placed medial to the SN for the VTA. A round shape ROI (1.6 × 1.6 mm^2^) was placed at the center of the caudate/putamen complex (ST) and frontal lobe (FL) region (Figure 1 D-F. The atlas template then was co-registered to the individual R2* maps using an affine registration with mutual information criteria using ANTs software to obtain the reference locations of the SN and other ROIs.^29^ The final ROIs were visually inspected to ensure location accuracy from registration step. For the lesioned rats in Experiment 1, the R2* values from both the lesioned and unlesioned side were analyzed. For the healthy rates in Experiment 2, the R2* values averaged from both sides were used for data analysis. The location and size of the SN also was verified using DAB-enhanced Perls’ staining (Figure 1B) and tyrosine hydroxylase (TH) staining (Figure 1C) from midbrain sections of one of the lesioned rats. The staining images demonstrated that the SN definition delineated by immunohistochemistry staining agreed with our definition on R2* images.

**Figure 1.**
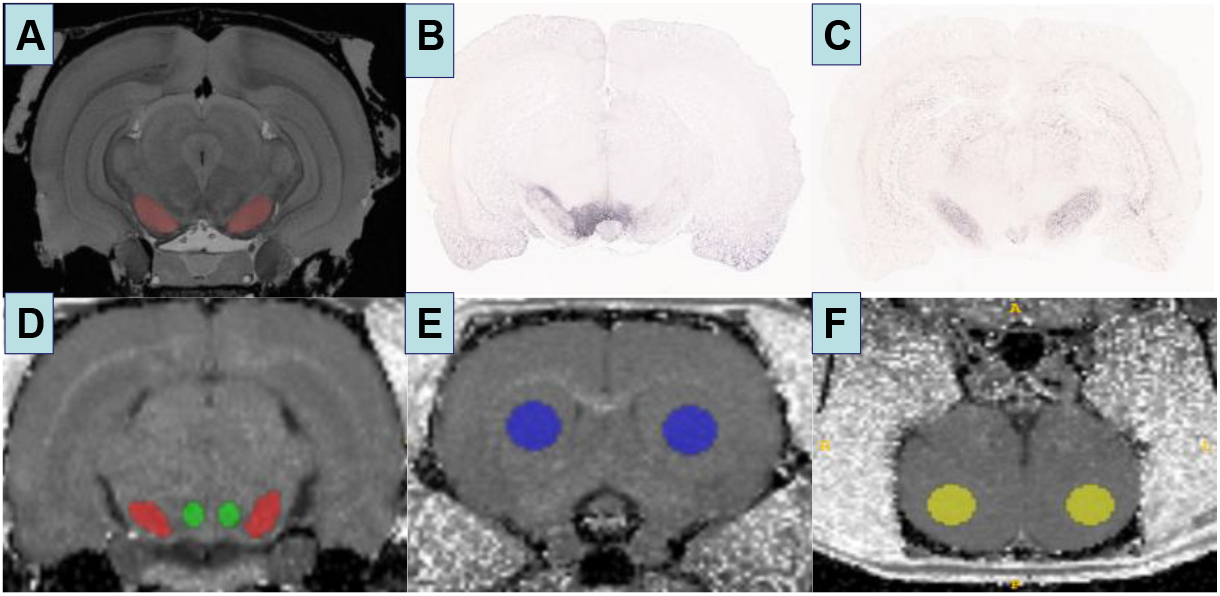
The SN is defined according to Waxholm Space Atlas of Rat Brain (A). The location and size of the SN also is validated from the same rat using TH staining (B) and Perls’ staining (C) images. An example case of the SN, VTA, ST, and FL definition in the R2* image from a lesioned rat is showed in (D-F).

### Statistical analysis

The primary outcome in Experiment 1 is the SN iron content change before and after levodopa treatment measured by the SN R2* values. Linear mixed models were used to assess R2* changes at the lesioned side, unlesioned side, and at the caudate/putamen complex respectively. A paired Student t-test provided a similar result. For Experiment 2, linear mixed models also were used to estimate the linear trend of iron change over 4 months with the SN R2* values as outcome and group, visit, and Group × Visit as independent variables. The Group × Visit term was used to assess the longitudinal effect of levodopa treatment on the SN iron content. We also used a Student’s *t* test to compare the baseline SN iron content from Experiment 1 and 2 before levodopa treatments to assess if there are any differences between 6-OHDA lesioned rats and healthy rats using.

## Results

### Levodopa increases iron in the SN in two different paradigms

The first experiment used the 6-OHDA unilaterally-lesioned model who were treated with levodopa for 15 days. The SN R2* values on the lesioned side increased from 22.41 before levodopa treatment to 23.08 after levodopa treatment (Figure 2; difference = 0.68, CI = 0.02 to 1.34, p = 0.042). On the intact side, the SN R2* values increased from 21.88 before levodopa treatment to 23.08 after levodopa treatment (difference = 1.19, CI = 0.43 to 1.95, p = 0.005).

**Figure 2.**
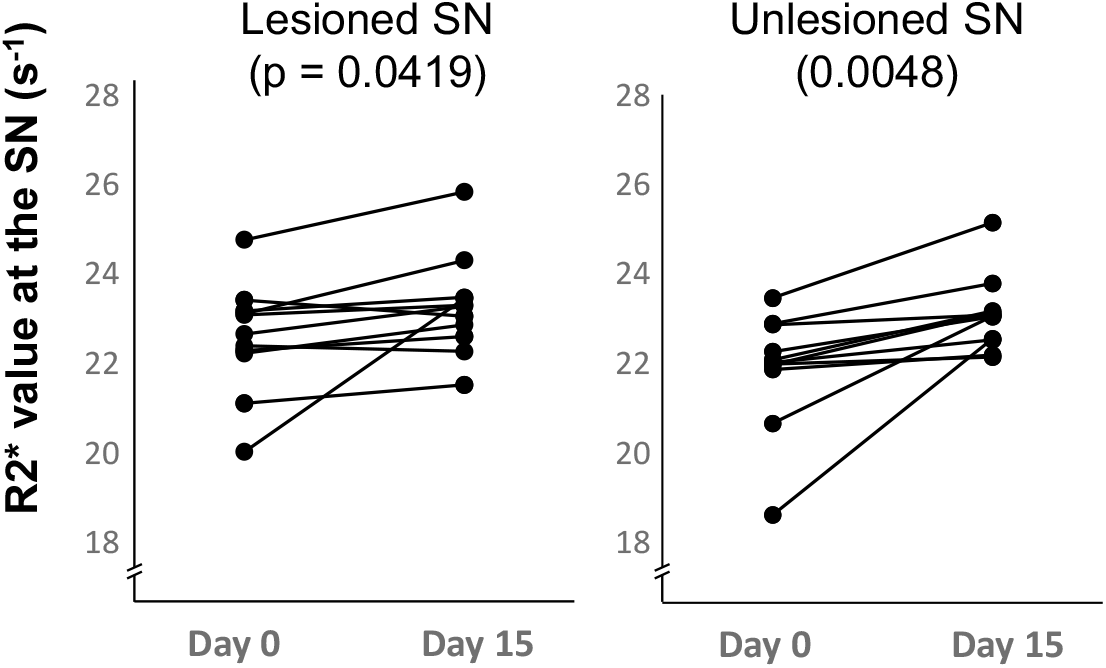
R2* changes before and after 15-day levodopa treatment on 6-OHDA PD rat model. The p-values are from mixed linear models without correcting for multiple comparisons.

The second experiment used unlesioned healthy rats. The levodopa treated group showed linear increase in iron content at the SN compared to the vehicle group [Figure 3 (Group × Visit term p = 0.013)]. The SN R2* values of the levodopa treated group increased from 21.12 at baseline to 22.53 in a 4-month period (difference = 1.42, CI = 0.49 to 2.34). Post-hoc tests showed that the levodopa treated group showed significantly higher SN R2* values at 2 months (p = 0.036) and at 4 months (p < 0.001) but not at 1 month (p = 0.575).

**Figure 3.**
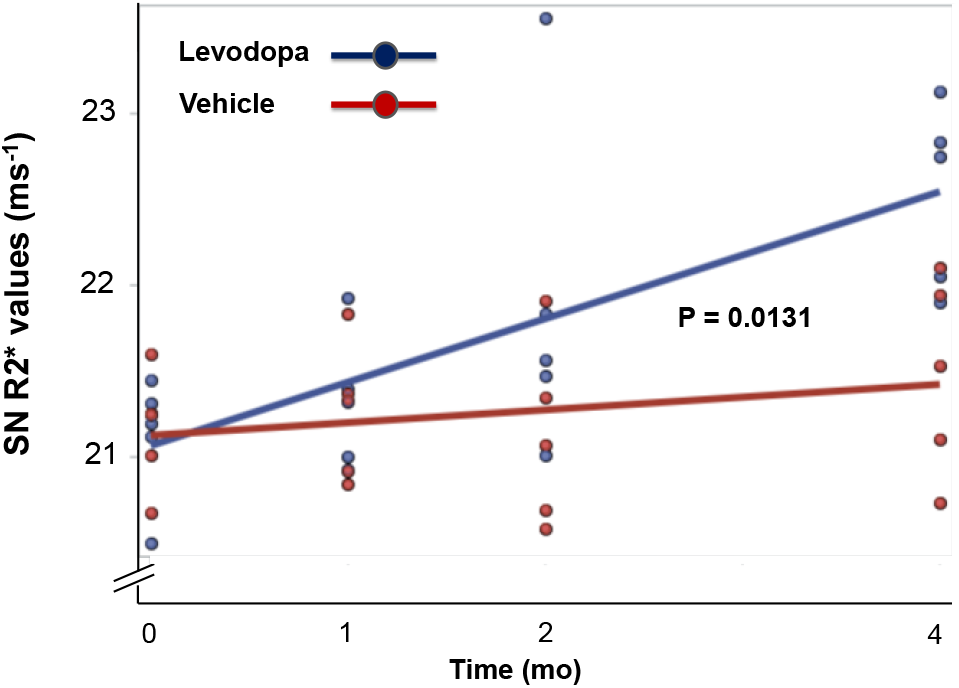
R2* changes after levodopa treatment of unlesioned rats for four months. The p-value is from linear mixed model indicating that the linear trajectory of treated group is significantly increased compared to vehicle group.

### Comparison of the SN iron content between 6-OHDA lesioned rats and healthy rats before levodopa treatment

The 6-OHDA-lesioned rats had higher iron content in the lesioned side compared to healthy rats [Lesioned side R2* mean (SD) = 22.40 (1.2), healthy rats R2* mean (SD) = 21.14 (0.34), p = 0.005] (Figure 4). The lesioned side also had a trend showing higher R2* values compared to the unlesioned side (p = 0.051). The unlesioned side had higher R2* values than that in 6-OHDA treated normal rats but this did not reach statistical significance [Unlesioned side R2* mean (SD) = 21.89 (1.30), healthy rats R2* mean (SD) = 21.14 (0.34), p = 0.093]. Of note, the SN R2* values in the 6-OHDA lesioned rats showed much greater variance than that seen in levodopa treated normal rats.

**Figure 4.**
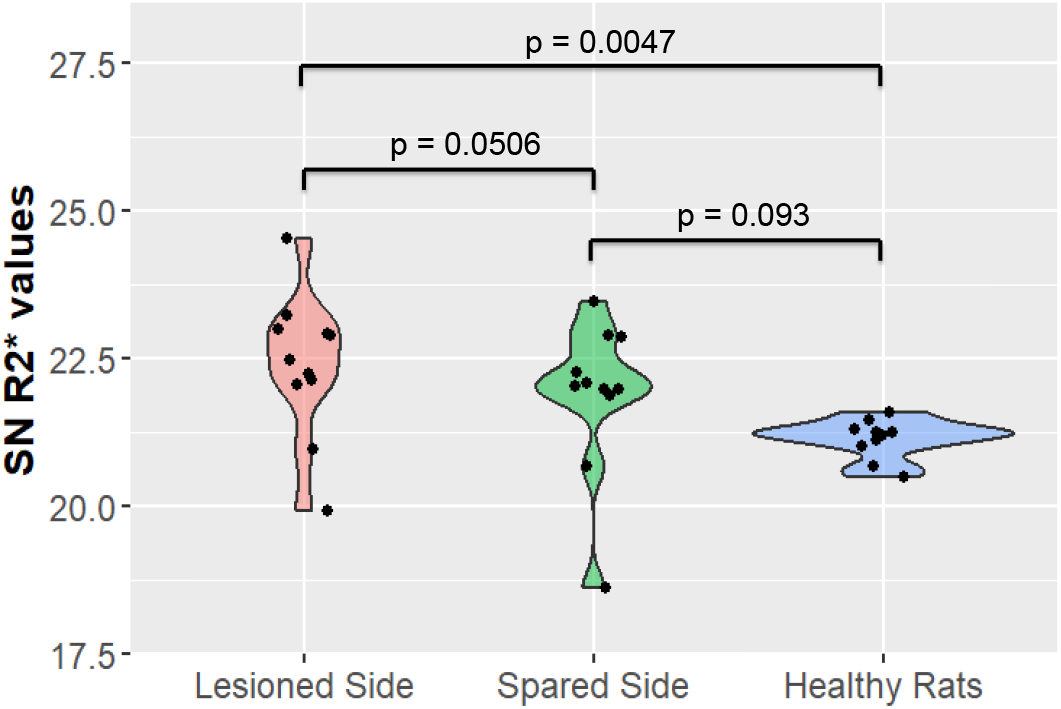
The baseline SN R2* values from Experiments 1 and 2 were compared to assess the SN iron change due to 6-OHDA lesioning. This was an exploratory analysis and the p-values (two-sided) from Student’s t tests, not corrected for multiple comparisons.

### Levodopa treatment does not cause increased iron levels in three dopamine terminal fields

Because of the marked changes in the SN, we examined whether levodopa would cause similar changes in several dopamine terminal fields. In the unilaterally lesion rats, there were no significant R2* changes observed in striatum, frontal lobe, or ventral tegmental area when comparing lesioned to unlesioned side at baseline or 15 days (see Figure 5). Conversely, the unlesioned side of VTA showed a trend of increase from baseline to follow-up (p = 0.0505), so this became another focus of experiment 2. The lesioned side, however, showed no R2* increase after levodopa treatment in the 15-day study.

**Figure 5.**
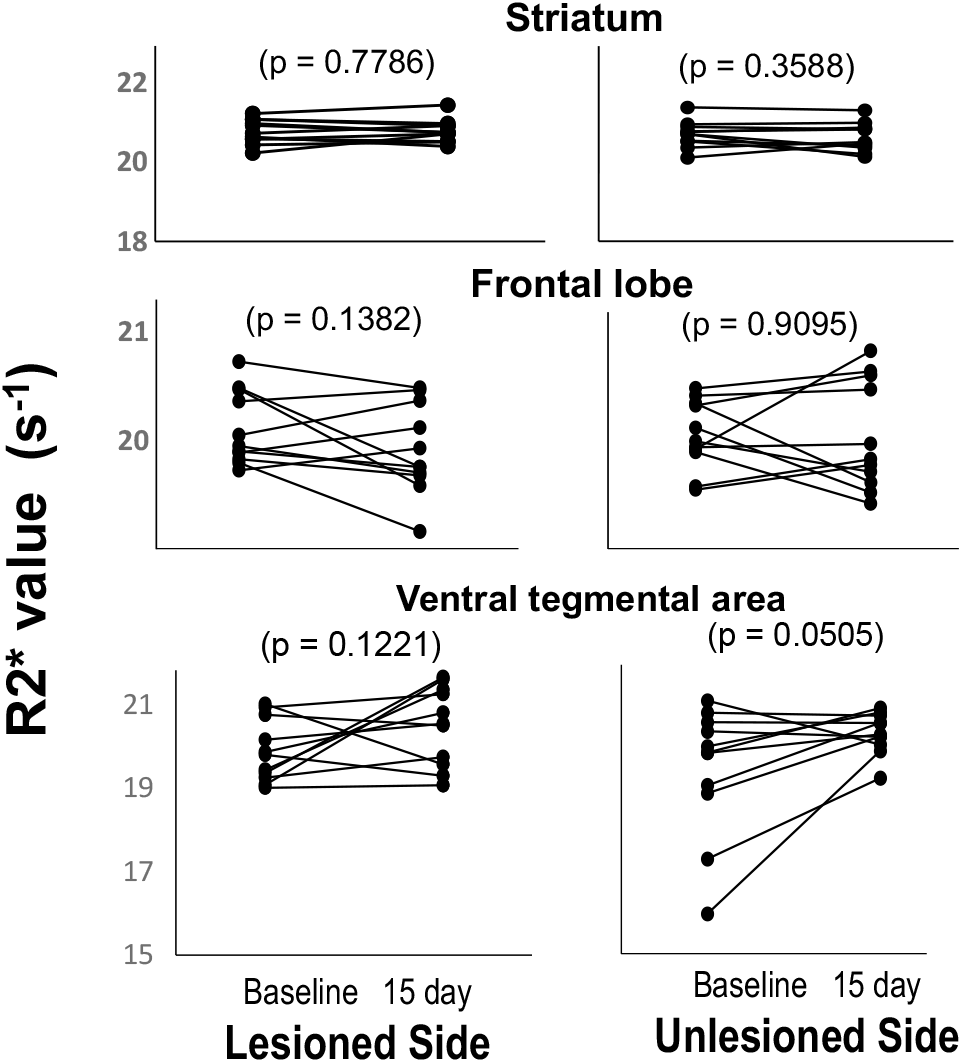
Effects of 15 days of levodopa in unilateral lesioned rats in the striatum, temporal lobe, and ventral tegmental area.

In experiment 2, unlike in the SN, levodopa did not cause any significant increases in R2* in striatum, frontal lobe, or ventral tegmental during the four-month epoch of the study (Figure 6). Although there was no significant effect of the levodopa treatment, it is interesting that there is a general trend towards higher iron concentrations as the animal age over the four-month epoch

**Figure 6.**
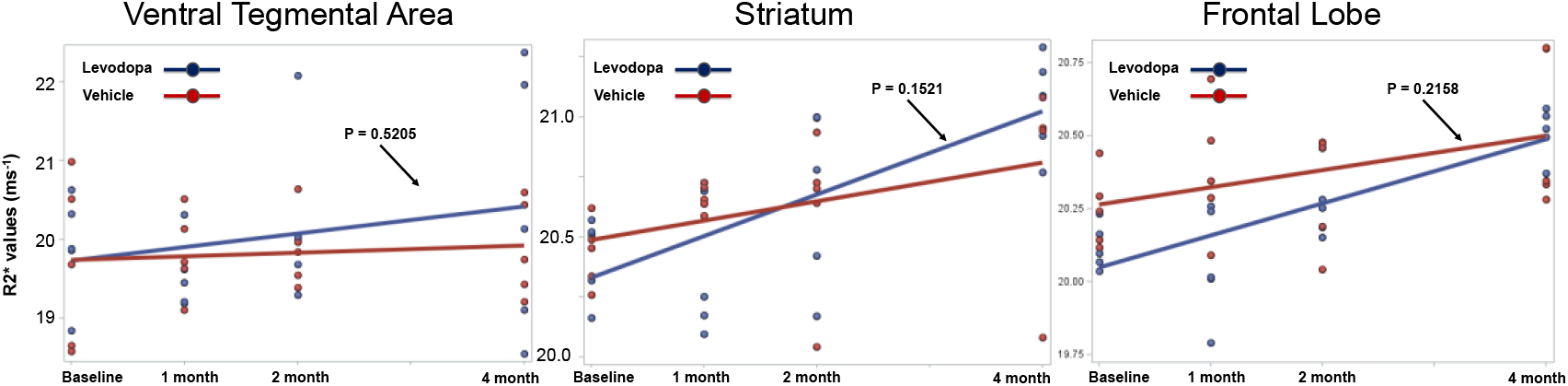
R2* changes after levodopa treatment of unlesioned rats for four months for the striatum, ventral tegmental area, and frontal lobe. The p-value is from linear mixed model indicating that the linear trajectory of treated group did not significantly increase compared to vehicle group.

## Discussion

Our original design principle was to use the unilateral 6-OHDA system not as a model of PD, but rather to take advantage of the lack of dopamine neurons on the lesioned side. Our premise was that damage to dopamine neurons from levodopa treatment results in an increase in iron (either as a response to, or mechanism of, injury). Thus, the lesioned side was predicted to have a limited capacity to respond because of its paucity (>90% loss) of dopamine neurons, whereas the unlesioned side should have a more robust response

Experiment 1 found that 15 days of levodopa treatment increased iron content consistent with the hypothesis formed from our previous MRI findings in PD patients.^21^ Yet contrary to the premise of that experiment, the levodopa treatment led to increased iron in <u>both</u> the lesioned and unlesioned SN. This was not predicted, and may represent the fact that there was activation of the immune system by 6-OHDA insult and that this played a role. It would be interesting to know if the post-lesioning period was markedly longer if a similar result would occur. Nonetheless, this led to a change in the design of experiment 2. We chose to use normal, unlesioned rats because it eliminated the secondary effects of responses related to the 6-OHDA insult (e.g., activation of the immune system). Experiment 2 found that levodopa treatment increased iron accumulation bilaterally in unlesioned animals.

Our analysis also found higher SN iron content in 6-OHDA-lesioned rats compared with healthy rats, consistent with prior studies.^30, 31^ This indicates there are independent effects of levodopa and of lesioning. Prior research using post-mortem histology has reported increased iron content in the SN after 6-OHDA lesioning and dopamine neuron degeneration,^31^ findings confirmed by other quantitative approaches.^30, 32^ Those studies, however, showed 20-30% increase in nigral iron in the lesioned side, but no significant increase on the unlesioned side. Our data result comparing baseline SN R2* values was consistent with these data in that only the lesioned side showed significant iron increase but not at the unlesioned side. Our results, however, showed increases in iron on both sides when the animals were treated with levodopa.

The exact clinical implication for levodopa-iron effects in healthy animal are unknown, as levodopa use is primarily for Parkinson’s disease and related parkinsonisms. Levodopa has, however, been used for restless leg syndrome. This practice has been discouraged due to intolerable augmentation effects, thus it is not a vehicle for a prospective study. There are a number of other non-neurodegenerative conditions such as levodopa-responsive dystonia, but the dosage of levodopa is lower than used in Parkinson’s patients. Levodopa has been used in a number of clinical conditions, both neurodegenerative (e.g., multiple system atrophy, progressive supranuclear palsy) and non-neurodegenerative (post-stroke). It will be useful to obtain baseline brain iron imaging and post-treatment imaging in those patients as a routine practice to determine iron responses to levodopa. In toto, iron-sensitive MRI can be very useful in studying these processes in vivo and in human, serving as a powerful and bidirectional translational tool.

### Levodopa administration and/or lesions increase iron in the substantia nigra

Whether levodopa has neurotoxic effects has been a point of controversy for decades, and the prevailing view is that levodopa is not neurotoxic or at least does not cause additional symptom progression. Earlier *in vitro* experiments showed that levodopa autoxidation causes oxidative stress, leading to neuronal death by necrosis or apoptosis. Chronic administration of levodopa to healthy rats or primates, however, generated inconclusive results. There is some evidence suggesting that, when the dopamine system is already damaged, levodopa treatment can produce further cell loss due to excessive oxidative stress. Our results from the 6-OHDA rat model are consistent with this notion. Levodopa treatment can lead to an increased iron level in both the lesioned side and unlesioned side. Despite TH positive neurons being well preserved in the unlesioned side, the unlesioned side can undergo compensatory expression upregulation of tyrosine hydroxylase and dopamine transporter.^33^ Due to the loss of dopamine neurons in the lesioned side, the unlesioned side might be responsible for an increased load of dopamine metabolism resulting in excessive iron uptake and deposition. The fact that nigral iron increases were observed in 15 days for the 6-OHDA lesioned rats compared to 2 months in healthy rats is consistent with the results from chronic administration of levodopa to healthy rats. Without preexisting stress, the dopamine neurons appear to tolerate levodopa treatment better. A recent *in vitro* study has demonstrated that levodopa treatment can lead to increased expression of an iron influx protein, divalent metal transporter-1 in glia.^34^ This might be one potential mechanism for levodopa inducing iron increases in the SN.

The most surprising result was that levodopa treatment can lead to an increase in nigral iron in healthy rats. This required a longer period time before the effect became statistically significant (2 months) compared to the 6-OHDA lesioned SN (15 days). If iron accumulation represents either a toxic mechanism or response to insult, it is clear that we have provided clear in vivo evidence for such a phenomenon. Most importantly, our data suggest that the impact of levodopa on nigral iron accumulation is chronic and independent of concomitant damage to dopamine neurons, suggesting parallel pathophysiological processes.

### Limitations

Our study has several limitations. The sample sizes for both experiments were not large, although the predicted effect size was large.^21^ The first experiment gave us confidence that we had adequate power, and the design was also helped by the fact that each animal also can be its own control, allowing us to use smaller numbers of animals. We also did not address in detail the temporal effects of the 6-OHDA lesioning process that tend to be dynamic for some months after lesion.^35-37^ This concern was obviated because our design was based on the fact that we were using each animal’s baseline as a control. In our study, the lesioned rats were given at least nine days of recovery to avoid the acute phase of dopamine neurons loss and possible residue effect of 6-OHDA on the SN iron content. Another limitation was the defining of the ROI for the SN that usually includes both the pars compacta and the pars reticulata. The spatial resolution limitation of MRI in the rat did not permit differentiating the role of glia residing in the pars reticulata of SN.

### Future directions

One of the largest past controversies in the PD field related to the idea that levodopa, despite its essential symptomatic effects, caused concomitant toxicity. This actually led to the standard-of-care for early PD shifting to one that was “levodopa sparing.”^18, 38-41^ The use of “dopamine agonists” (i.e., D_2_-selective drugs) in place of levodopa in early disease was designed to delay “toxic” effects of levodopa until patients could not tolerate consequent inadequate symptom control. In 1996, Fahn^42^ wrote: “*The long-running controversy whether levodopa is the culprit or the bystander in producing motor response fluctuations and the more recent concern as to whether it can hasten the progression of PD by producing increasing oxidative stress need [sic] to be answered; a prospective, controlled clinical trial might be able to provide these answers*.*”* This laid the foundation for two landmark studies: ELLDOPA^19^ and LEAP.^43^ When these studies showed functional benefit in the levodopa-treated subjects,^*44, 45*^ it was considered the “*Final Nail in the Coffin of Disease Modification for Dopaminergic Therapies*.”^46, 47^

Our data suggest that the “coffin” should be opened again with 21^st^-century new toolsets. Basic biology would predict that by increasing dopamine neurotransmission, levodopa use in early and mid-stage disease will normalize brain circuitry, improving brain health. This would explain the beneficial effects seen in ELLDOPA. But if nigral iron is a marker or mediator of brain toxicity, it also suggests that levodopa could be causing covert damage whose effects cannot be detected because any detrimental effects are masked by the essential symptomatic effects of levodopa. This could explain the decreases in imaging biomarkers of dopamine terminal integrity. Unfortunately, neither ELLDOPA nor LEAP could deliver an unambiguous answer because there was no true control group (i.e., a treatment of equal efficacy to levodopa but without the chemical features of levodopa presumed to mediate toxicity).

As noted earlier, we found that increased SN iron did not occur in PD patients until after diagnosis,^21^ concomitant with initiation of levodopa therapy. The prior human study, however, was cross-sectional in nature could not prove that the findings were not a consequence of the disease alone. The current study now provides direct evidence that these effects seen in patients may be explained by the actions of levodopa. There are two directions that can answer this question. The first is to reexamine the mechanisms that might lead to iron accumulation. Since iron accumulation occurs even on the 6-OHDA lesioned side, it suggests cell types other than dopamine neurons are involved (e.g., microglia or astrocytes).^48-50^ The accelerated effect in animals that had been lesioned with 6-OHDA also suggest that neuronal insult (as occurs in PD) might synergize these actions of levodopa.

Proving these mechanisms is now possible clinically with advances in MRI imaging. Moreover, what was missing in ELLDOPA and LEAP was a proper reference group (i.e., a comparator drug of equal antiparkinson efficacy but devoid of the reactivity of levodopa). In this regard, decades of research have shown that one class of drug, selective dopamine D_1_ agonists, might equal levodopa in efficacy.^51-55^ The pharmaceutical limitations of early D_1_ agonist drug candidates have been recently overcome. Phase 3 data for tavapadon, an orally-available long-acting partial D_1_ agonist, are now available and suggest equal efficacy to levodopa.^56, 57^ Tavapadon does not contain the catechol-chemical moiety that is proposed to be the reactive part of levodopa.^58^ Because the Phase 3 data suggests that tavapadon can be effective monotherapy, it now could provide a useful reference population. The current data would predict that iron accumulation in PD patients would be markedly lower in the tavapadon-treated patients versus those taking levodopa. A corollary is the prediction that this will slow disease progression, at least that part that now can be associated with levodopa. This should lead to measurable functional benefits, and has transformative implications for Parkinson’s patients and related disorders and neuroscience community

## Acknowledgments

The authors thank Susan Kocher and Natalia Loktionova for their invaluable technical support. This work was supported by the National Institutes of Health (R01 AG071675, U01 NS112008, R01 ES019672, U01 NS082151, R01 NS060722), the Glatfelter Family and Michael J Fox Foundations, the Children’s Miracle Network, the Penn State Translational Brain Research Center, a Penn State Center for Biodevices Seed Grant, and University of Virginia research development funds.

